# Transient lagging chromosomes cause primary microcephaly

**DOI:** 10.1101/2024.05.02.592199

**Authors:** Elena Doria, Daria Ivanova, Alexandre Thomas, Patrick Meraldi

**Affiliations:** Department of Cell Physiology and Metabolism, Faculty of Medicine, University of Geneva, 1211 Geneva 4, Switzerland; Translational Research Centre in Onco-hematology, Faculty of Medicine, University of Geneva, 1211 Geneva 4, Switzerland

## Abstract

Primary microcephaly results from the impaired neuronal progenitor proliferation, causing reduced brain size and impaired cognitive abilities. Loss of the most frequently microcephaly genes, *WDR62* and *ASPM*, slows down poleward microtubule flux and results in transient lagging chromosomes in anaphase. Whether these defects cause primary microcephaly is unknown. Here we show that transient lagging chromosomes elicit an Aurora-B-dependent activation of 53BP1 and p21, impairing cell proliferation. Co-depletion of the microtubule depolymerase inhibitor CAMSAP1/Patronin in a WDR62 depletion background, restores normal flux rates, suppresses lagging chromosomes, and allows normal cell proliferation in human cells, while rescuing the small brain and the cognitive defects in fly larvae. We postulate that transient lagging chromosomes in anaphase and 53BP1/p21-dependent response they elicit, are a major driver of primary microcephaly.

## INTRODUCTION

Autosomal recessive primary microcephaly (microcephaly primary hereditary [MCPH]) leads to smaller brain size in new-borns due to exhaustion of neuronal progenitor cells (*1*). Recessive loss-of-function mutations in 30 different genes, most of which involved in cell cycle and cell division, are linked to this disease. This includes genes encoding key regulators of centriole biogenesis (*PLK4, SAS6, STIL, SAS4, CEP152* and *CEP135),* pericentriolar proteins (*CDK5RAP2, PCNT, TUBGCP4* and TUBGCP6), microtubule-associated proteins and microtubule-motors of the mitotic spindle (*ASPM, WDR62, MAP11*, *KIF11* and *KIF14*), kinetochore proteins (*KNL1* and *CENP-E),* and regulators of DNA damage/condensation/replication (*MCPH1, ORC1, ORC4, ORC6, CDC45, NCAPH, NCAPD2* and *NCAPD3)* (*2–5*). The common denominator between those genes remains unknown. For the cell division genes, several non-exclusive hypotheses have been postulated: (1) chromosome segregation errors that lead to aneuploidy causing neuronal progenitor pool depletion via p53-induced apoptosis (*6*); (2) erroneous spindle orientation that drives premature neuronal differentiation depleting the neuronal progenitor pool (*7–9*), (3) or centrosomal defects that prolong mitosis, causing an activation of 53BP1 that triggers a p53-dependent cell proliferation arrest (*10*). However, loss of function of WD Repeat-Containing Protein 62 (*WDR62*, MCPH2) and the abnormal spindle-like microcephaly associated gene (*ASPM*, MCPH5), which account for over 80% of all microcephaly cases (*11*), have not been conclusively linked to aneuploidy, spindle mis-orientation or a mitotic delay in human cells (*12–14*).

Instead, WDR62 and ASPM regulate microtubule dynamics by recruiting the microtubule-severing enzyme Katanin to mitotic spindle poles (*14–16*). Impairment of either protein results in insufficient microtubule minus-end depolymerization and reduced poleward microtubule flux rates (*14–16*), a conveyor belt-like movement that drives tubulin subunits towards the minus ends at spindle poles (*17*). We previously showed that human WDR62 depletion does not delay mitosis but instead leads to transient lagging chromosomes in anaphase that are nevertheless incorporated into the daughter nuclei (*14*). Whether these transient chromosome segregation defects are linked to microcephaly remains unknown.

Here, we show that transient lagging chromosomes in anaphase in WDR62- and ASPM-depleted human cells elicit an Aurora-B dependent response that activates 53BP1 and delays cell proliferation. We hypothesize that such mild segregation defects, which are frequently seen after the loss of most MCPH genes, could be a common origin for the reduced proliferation of neuronal progenitor cells. To test this hypothesis, we co-depleted CAMSAP1, a member of the Patronin/CAMSAP family, which inhibits spindle-associated microtubule depolymerases of the kinesin-13 family at spindle poles (*18–22*). CAMSAP1/Patronin co-depletion in a WDR62 RNAi background suppressed the appearance of transient lagging chromosomes and the cell proliferation delay in human cells and *Drosophila melanogaster* neuroblasts, and rescued microcephaly in fly larvae. We postulate that transient lagging chromosomes in anaphase are a major cause of primary microcephaly.

## RESULTS

Depletion or deletion of WDR62 in human cells leads to decreased microtubule depolymerization at spindle poles, slower poleward microtubule flux rates, and transient lagging chromosomes in anaphase (*14*). Such chromosomes (also called lazy chromosomes) transiently lag behind the DNA masses during anaphase in WDR62-depleted cells but, unlike persistent lagging chromosomes, are incorporated into daughter nuclei without formation of micronuclei (*14*, *23*). Transient lagging anaphase chromosomes were visible in live WDR62-depleted non-transformed retina pigment epithelial cells expressing human telomerase (hTert-RPE1) stained with SiR-DNA (p = 0.0191 in paired t-test; Figure 1A and B and Supplementary Figure 1A and B for siRNA validation). These lagging chromosomes reflected asynchronous kinetochore movements, since such laggards were also visible in WDR62-depleted RPE1 cells expressing kinetochore (GFP-CENP-A) and centrosome (GFP-Centrin1) markers (p = 0.0087 in paired t-test; Supplementary Figure 1C and D). Transient lagging chromosomes had so far not thought to lead to DNA damage. Since primary microcephaly is genetically also linked to DNA damage signalling, we nevertheless tested whether WDR62 depletion activated such pathways, by staining for γH2AX and 53BP1, and using the DNA-damaging agent Doxorubicin as positive control. Both proteins form nuclear foci upon activation: while γH2AX labels double-strand breaks (*24*), 53BP1 is involved in double-strand break repair, but also in p53 stabilization after long mitotic delays (*25–29*). This drives the expression of the cyclin-dependent kinase inhibitor p21, a key p53 target, resulting in cell cycle delay or arrest (*30*). Using automatic image analyses (Supplementary Figure 1E) on hundreds of interphase cells per experiment, we found that WDR62 depletion increased the proportion of cells with 53BP1-positive nuclei in each independent experiment, from an average of 31% (siControl) to 42% (siWDR62; overall p < 0.0001 in Fischer’s exact test, Fig. 1C and D), but did not increase the proportion of γH2AX-positive cells (Supplementary Figure 1F and G). WDR62-depleted cells typically displayed single nuclear 53BP1-positive foci, while doxorubicin-treated cells displayed dozens of 53BP1 foci (Figure 1C). The same automated analysis also indicated that WDR62-depleted cells had elevated nuclear p21 levels (p < 0.0001 in Fisher’s exact test; Figure 1C and E). When we labelled control- or WDR62-depleted cells with cell proliferation pulse-chase marker CFSE (Carboxyfluorescein succinimidyl ester, Supplementary Figure 1H for methodology) and let them proliferate for 5 days, we found that WDR62-depleted cells underwent half a cell division less than control-depleted cells (p = 0.0029 in one way ANOVA; in comparison serum starved cells underwent 2 cell division less; Figure 1F and G). This implied that WDR62-depletion activates the 53BP1-p53-p21 signalling pathway and delays cell proliferation independently of double-strand breaks. Consistently, WDR62 depletion did not significantly delay cell proliferation in a p21 knockout RPE1 cell line, while delaying cell proliferation in the parental cell line (p = 0.0141 in one way ANOVA; Figure 1H).

**Figure 1:**
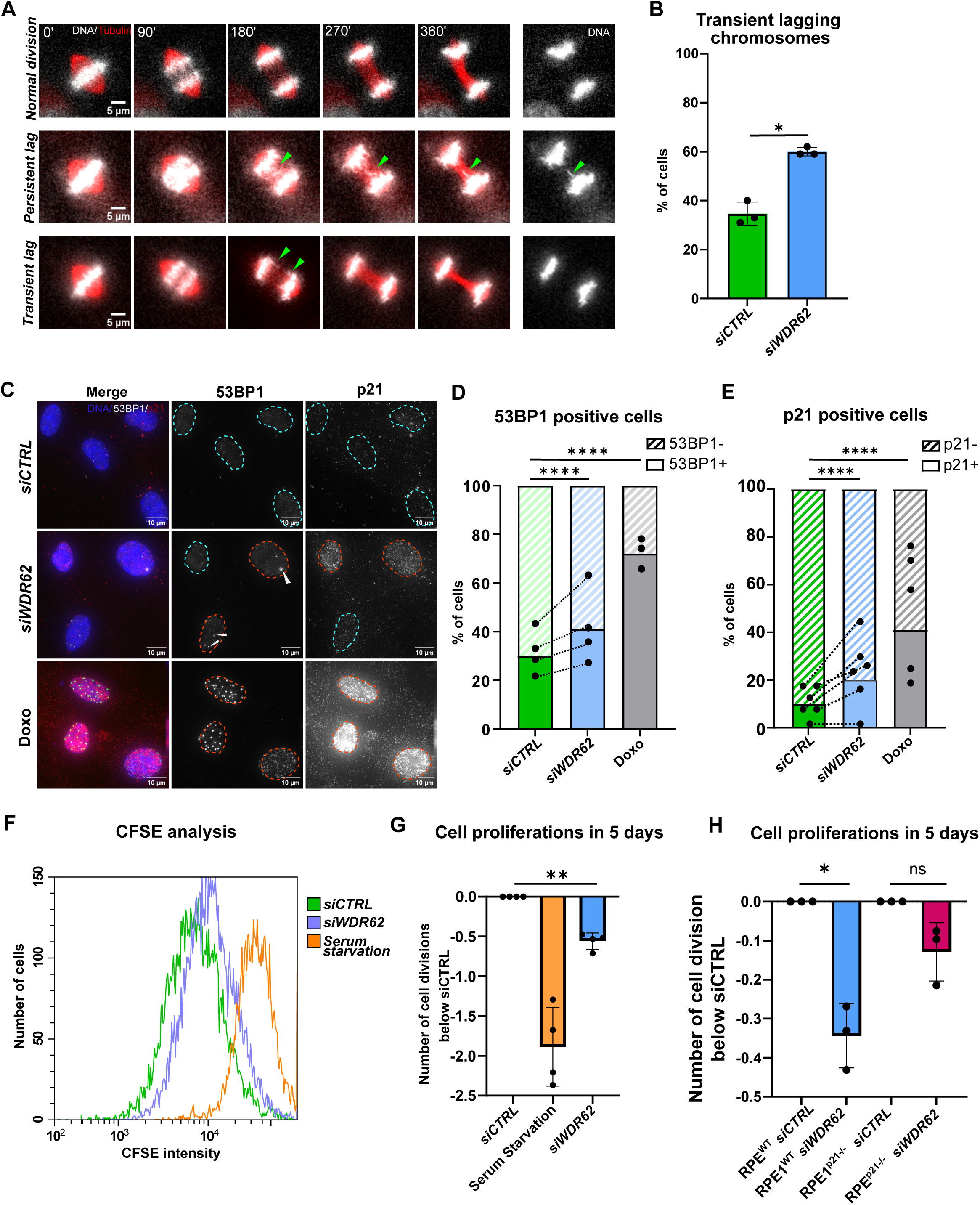
WDR62 depletion leads to 53BP1 and p21 activation and a proliferation delay. (**A**) Time-lapse single-plane images RPE1 cells stained with SiR-Tubulin (red) and SpY-DNA (white) showing examples of transient or persistent lagging chromosomes (green arrows). Scale bars = 5 µm. (**B**) Quantification of transient lagging chromosomes in *siWDR62-*treated RPE1. A cell was considered to have a transient lagging chromosome if a chromosome arm was sticking out of the DNA mass by >2 μm as the two DNA masses were separated by >6 μm; N = 3 independent experiments; n = 83-115 cells, *p = 0.0191, student two-tailed t-test. (**C**) Immunofluorescence images of interphase RPE1 cells treated with *siCTRL*, *siWDR62* or Doxorubicin, and stained with DAPI, 53BP1 and p21 antibodies. Blue dotted lines represent 53BP1-negative, red ones 53BP1-positive nuclei, arrow indicate 53BP1-foci in WDR62-depleted cells. Scale bars = 10 µm. (**D**) Quantification of 53BP1 positive cells. Connected dots represent paired experiments. N = 4 independent experiments, n *=* 648 – 1440 cells; ****p < 0.0001, Fisher’s exact test with Bonferroni correction (**E**) Quantification of p21 positive cells. Connected dots represent paired experiments. Blue dotted lines represent p21-negative, red ones p21-positive nuclei. N = 6, n = 570-1010 cells; ****p < 0.0001, Fisher’s exact test with Bonferroni correction (**F**) Example of FACS CFSE analysis data (**G**) Quantification of cell divisions after 5 days compared to control depletion based on CFSE intensity, N = 4, *n* = 18194 - 82963 cells; **p = 0.0029, one-way ANOVA test. (**H**) Quantification of cell divisions after 5 days when compared to control depletion based on CFSE intensity in RPE1 WT or p21^-/-^ cells, N = 3, *n* = 27693 - 29379 cells; *p = 0.0141, one-way ANOVA test. All error bars represent SEM.

### Transient lagging chromosomes induce 53BP1 foci and delay the cell cycle via Aurora B signalling

53BP1 is activated in the absence of DNA damage, but also if cells are delayed in mitosis, for example after centrosome loss (*27–29*). Aneuploidy can also activate p53 (*31*). WDR62 depletion, however, does not lead to a mitotic delay or micronuclei, nor does it affect centrosome numbers (*14*). Metaphase spreads over 40 cells in 3 replicates indicated that WDR62-depleted RPE1 cells had generally the same chromosome numbers as control-depleted cells; in contrast cells treated with Reversine, an inhibitor of the spindle assembly checkpoint kinase Mps1 that induces aneuploidy, displayed severe aneuploidy ((*32*); Supplementary Figure 2A and B). To exclude low levels of aneuploidy and quantify individual chromosomes at single cell level, we compared control- and WDR62-depleted RPE1 cells by single cell DNA sequencing. We found that within a population of 384 cells, WDR62- and control-depleted cells had the same chromosome distribution (Figure 2A, and supplementary Figure 2C), confirming that WDR62 depletion does not induce aneuploidy. We next reasoned that 53BP1 activation could be either linked to the transient lagging chromosomes or to an unrelated cell-cycle function of WDR62. To differentiate the two possibilities, we tested whether the novel appearance of 53BP1 foci correlated with the presence of transient lagging chromosomes in the preceding anaphase. Monitoring live WDR62-depleted 53BP1-GFP RPE1 cells labelled with SiR-DNA (*33*), revealed that the appearance of 53BP1 nuclear foci in hitherto 53BP1-negative G1 cells was nearly always preceded by a transient lagging chromosome in anaphase (p = 0.0105 in student t-test; Figure 2B and C). This suggested that transient lagging chromosomes are linked to 53BP1 foci formation. Consistently, ASPM-depleted cells also displayed transient lagging chromosomes (p = 0.0005 in paired t-test) and 53BP1 foci (p < 0.0001 in Fischer’s test with Bonferroni correction; Figure 2D-G, and supplementary Figure 2D and E for siRNA validation).

**Figure 2:**
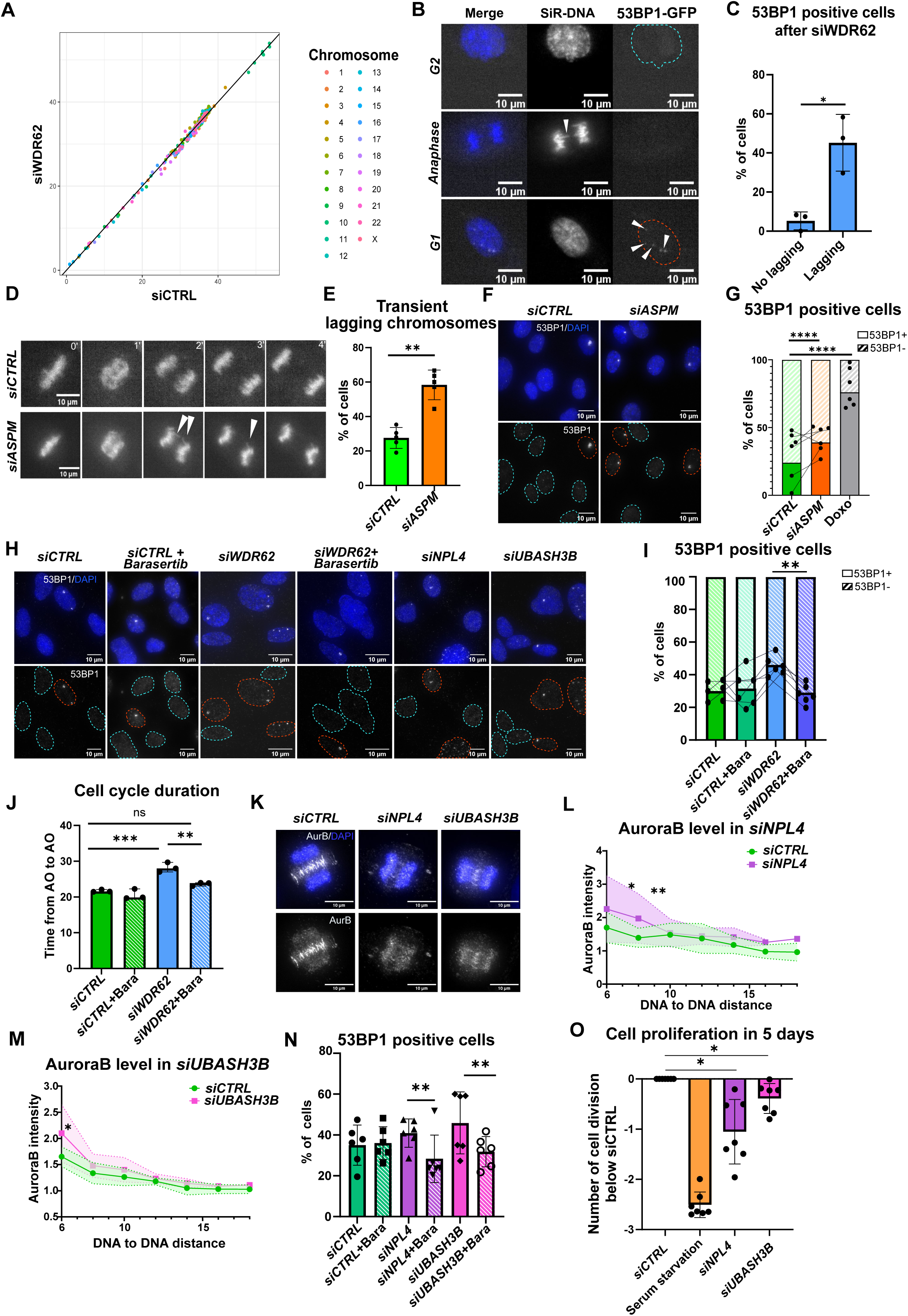
Transient lagging chromosomes activate 53BP1 via Aurora B. **(A)** Scatter-plot of chromosome copy number comparing the coverage of individual chromosomes in WDR62-vs. control-depleted cells. Numbers indicate the normalized number of reads per chromosome. The diagonal indicates that the chromosome numbers in each dataset are highly correlated **(B)** Time lapse images of RPE-1 53BP1-GFP cell treated with *siWDR62* in G2 (top row, blue dotted circle 53BP1-negative), anaphase (middle row) and in the ensuing G1 phase (bottom row, red dotted circle 53BP1-positive). White arrow indicates transient lagging chromosome. Scale bars = 10 µm **(C)** Quantification of *siWDR62*-treated 53BP1-GTP positive cells after a cell division with or without lagging chromosomes. N = 3; n = 60 cells; *p = 0.0105, student two-tailed t-test. (**D**) Time lapse imaging of RPE1 cells stained with SiR-DNA undergoing mitosis. Arrows indicate transient lagging chromosomes. Scale bars = 10 µm **(E)** Quantification of lagging chromosome in SiR-DNA stained live RPE-1 cells treated with *siCTRL* or *siASPM*. N = 5; n = 41-115 cells; ***p = 0.0005, paired t-test. **(F)** Immunofluorescence images of siCTRL- or siASPM-treated RPE-1 cells stained for 53BP1 and DAPI. Blue dotted circles represent 53BP1-negative, red ones 53BP1-positive nuclei. **(G)** Quantification of 53BP1 positive cells after *siCTRL* (n=1556 cells), *siASPM* (n=1224 cells) or Doxorubicine (n=681 cells) treatment. N = 6, n = 681-1556 cells; connected dots represent paired experiment. ****p < 0.0001, Fisher’s exact test with Bonferroni correction. **(H)** Immunofluorescence images of RPE-1 cells treated with indicated treatments and stained for 53BP1 and DAPI. Light blue dotted circles represent 53BP1-nuclei, red ones 53BP1 positive nuclei. Scale bars = 10 µm **(I)** Quantification of 53BP1 positive cells after *siCTRL*, *siCTRL* + barasertib, *siWDR2*, *siWDR62* + barasertib treatments. N = 6, n = 1516-1883 cells; connected dots represent paired experiments. *siCTRL vs siWDR62* p = 0.0082, *siWDR62 vs siWDR62 +* barasertib p = 0.0051, one-way Anova test. (**J**) Quantification of cell cycle duration (anaphase to anaphase) after indicated treatments. N = 3, n = 103 – 129 cells. *siCTRL* vs *siWDR62* p = 0.0134, *siWDR62* vs *siWDR62* + barasertib p = 0.0085, one-way Anova test. (**K**) Immunofluorescence of RPE-1 cells after *siCTRL*, *siNPL4* or *siUBASH3B* treatment stained for AuroraB and DAPI. Scale bars = 5 µm **(L and M)** Aurora B intensity quantification in anaphase RPE-1 cells treated with *siCTRL*, *siNPL4* or *siUBASH3B*. Y-axes indicate Aurora B intensity on DNA normalized to background; x-axes indicates the distance between the two DNA masses reflecting anaphase progression, dotted lines represent 95% confidence intervalls, n = 77 -113 cells; *p = 0.0369, **p = 0.0034 (L) and *p = 0.0400 (M), Mann-Whitney test **(N)** Quantification of 53BP1 positive cells after indicated treatment. N=8, n = 1900-2790 cells, *siCTRL* vs *siNPL4* p = 0.0094, *siCTRL* vs *siUBASH3B* p = 0.0274, one-way Anova test **(O)** Quantification of cell divisions after 5 days when compared to control depletion based on CFSE intensity in NPL4- and UBASH3B-depleted cells, N = 7, *n* = 36’799-49’501 cells; *siCTRL* vs *siNPL4* p = 0.121 *siCTRL* vs *siUBASH3B* p = 0.0326, one-way Anova test. All error bars represent SEM.

Why would transient lagging chromosomes in anaphase activate 53BP1? One possibility is that transient lagging chromosomes are exposed longer to the activity gradient of the mitotic kinase Aurora B at the spindle mid-zone (*34*). Indeed, Aurora B activity is high at the spindle midzone, where it can phosphorylate chromosomal components if chromosomes are lagging, and low in the regions of spindle poles (*34*). The Aurora B gradient can delay nuclear envelope reformation (*35*, *36*), and has been proposed to activate p53 in the presence of permanent lagging chromosomes (*37*). If correct, partial Aurora B depletion should prevent 53BP1 foci formation and the cell cycle delay in WDR62-depleted cells. Partial inhibition of Aurora B with 50 nM Barasertib, a condition that did not prevent cytokinesis but reduced the Aurora B-dependent localization of MCAK at kinetochores by 60% ((*38*) Supplementary Figure 2F and G), indeed abolished the increase in 53BP1-positive nuclei in WDR62-depleted cells (Fig. 2H and I). Moreover, when we extracted cell cycle duration (time between two mitoses) from long-term live cell imaging we found that partial Aurora B inhibition suppressed the 9 hour cell cycle delay induced by WDR62 depletion (p = 0.0115 in one-way Anova test, Figure 2J and supplementary Figure 2H and I). This indicated that the 53BP1 foci formation and the cell cycle delay in WDR62-depleted cells depend on Aurora B activity.

Aurora B is extracted from chromosomes at anaphase onset via Cdc48/p97 and the de-ubiquitinase UBASH3B, allowing its transport to the spindle midzone (*36*, *39*, *40*). We therefore predicted that maintaining Aurora B on chromosomes should lead to more 53BP1-positive cells and a reduced proliferation. Partial depletion of the Cdc48/p97 co-factors NPL4 or UBASH3B, delayed chromosomal Aurora-B extraction in anaphase (p = 0.040 and p = 0.0034 and in Mann-Whitney test, Figure 2K-M, Supplementary Figure 2J-M), and led indeed to more 53BP1-positive cells (Figure 2H - N) and a reduced cell proliferation (Figure 2O), despite the absence of a mitotic delay or lagging chromosomes (Supplementary Figure 2N-P and data not shown). Consistent with our model, the increase in 53BP1 positive cells after NPL4 or UBASH3B depletion was suppressed by partial Aurora B inhibition (Figure 2N). Overall, this implied that a deregulation of poleward microtubule flux in WDR62-depleted cells favours the transient presence of lagging chromosomes in a high Aurora B activity gradient, causing 53BP1 activation and a cell cycle delay.

### Co-depletion of CAMSAP1 rescues poleward microtubule flux rates and lagging chromosomes in WDR62-depleted cells

To test this hypothesis, we aimed to restore normal poleward microtubule flux rates in WDR62-depleted cells. One promising candidate was the depletion of CAMSAP1, a member of the CAMSAP protein family, which we found to localize to spindle poles in mitotic cells (Figure 3A and B). CAMSAP1-3 proteins and their *Drosophila melanogaster* ortholog Patronin inhibit the microtubule depolymerizing activity of kinesin-13s; moreover, *PATRONIN* mutations increase poleward microtubule flux rates in flies (*18–20*). In RPE1 cells, CAMSAP1 depletion increased poleward microtubule flux rates in metaphase by 50% and doubled the percentage of cells displaying transient lagging chromosomes (Figure 3C-F). This suggested that both a reduction and an increase in poleward microtubule flux impair efficient anaphase chromosome movements. Co-depletion of CAMSAP1 and WDR62, however, restored normal poleward microtubule flux rates (Figure 3G-K) and rescued the transient lagging chromosome phenotype (Figure 3L). We conclude that the reduction of poleward microtubule flux causes lagging chromosomes in WDR62-depleted cells, and that efficient anaphase chromosome movements require a balanced microtubule flux rate.

**Figure 3:**
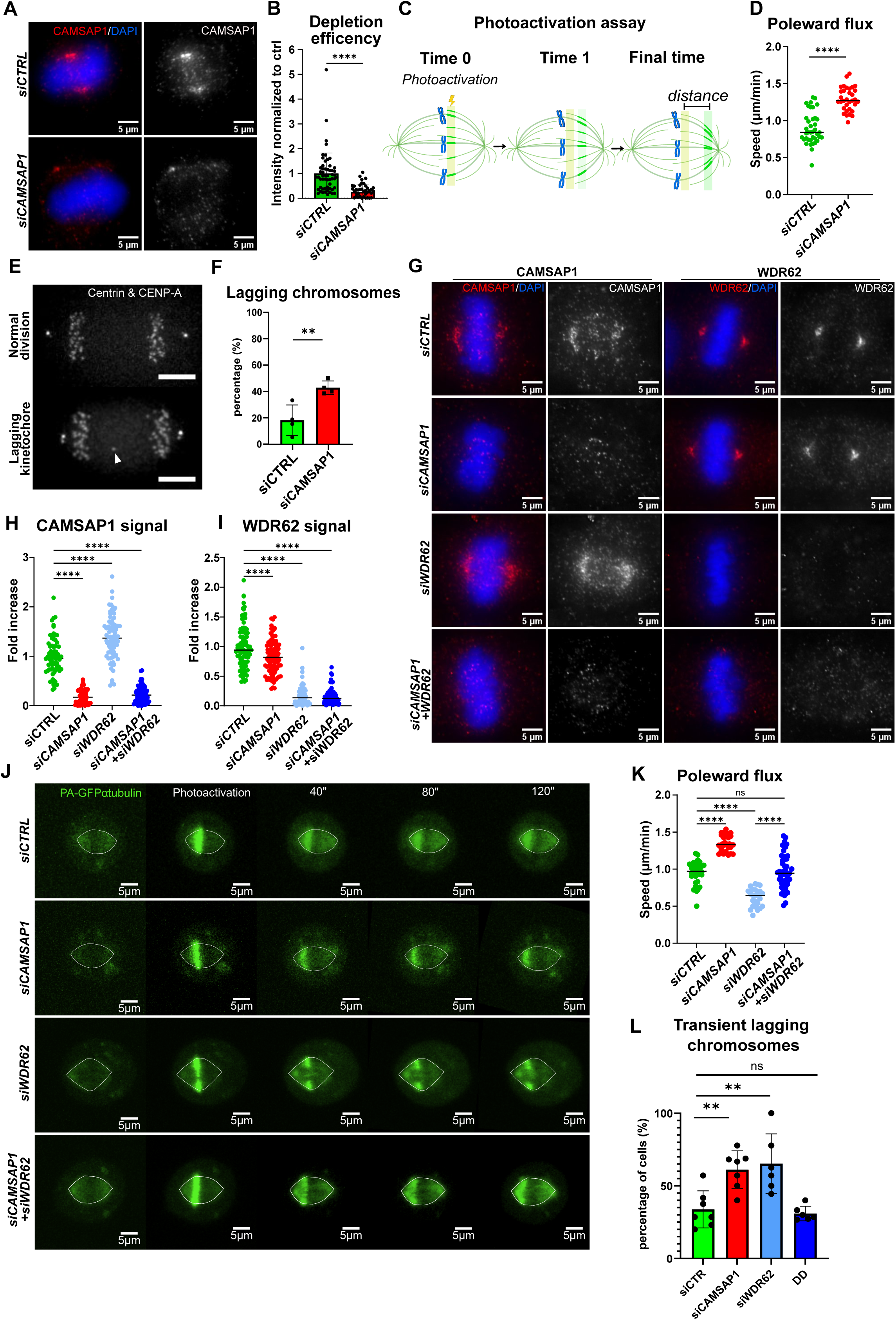
CAMSAP1 co-depletion rescues the transient lagging chromosomes in WDR62-depleted cells. **(A)** Immunofluorescence of RPE-1 cells treated with *siCTRL* or *siCAMSAP1* stained for CAMSAP1 and DAPI. **(B)** Quantification of normalized CAMSAP1 level in RPE-1 cells treated with *siCTRL* (n=55 cells) and *siCAMSAP1* (n=47 cells). N = 3, n = 47-55 cells, p = 0.0103, student two-tailed t-test. **(C)** Scheme of photoactivation assay for microtubules poleward flux measurement. **(D)** Quantification of microtubule poleward flux speed in RPE-1 PAGFP-αtubulin cells treated with *siCTRL* and *siCAMSAP1*. N= 3, n = 33-39 cells, p = 0.0008, paired t-test. **(E)** Stills of RPE-1 GFP-Centrin1/GFP-CENPA cells in a normal anaphase or an anaphase with a transient lagging kinetochore. **(F)** Quantification of transient lagging chromosomes in 15 minutes live cell movies of RPE1 GFP-Centrin1/GFP-CENPA cells treated with *siCTRL* or *siCAMSAP1*, n = 66-71 cells, p = 0.0071, paired t-test. **(G)** Immunofluorescence images of RPE-1 cells treated with *siCTRL*, *siCAMSAP1*, *siWDR62*, or *siCAMSAP1 + siWDR62,* stained for CAMSAP1 (left panels), WDR62 (right panels) and DAPI. **(H)** Quantification of normalized CAMSAP1 levels in RPE-1 cells treated with indicated siRNAs. N=3, n = 66-95 cells, *siCTRL vs siCAMSAP1* p = 0.0008, *siCTRL vs siCAMSAP1 + siWDR62* p = 0.0013 in one-way Anova test. **(I)** Quantification of normalized WDR62 levels in RPE-1 cells treated with indicated siRNAs. N=3, n = 66-95 cells, *siCTRL vs siCAMSAP1* ****p < 0.0001 in one-way Anova test. **(J)** Live cell images RPE-1 PAGFP-αtubulin cells treated with indicated siRNAs before and after photoactivation **(K)** Quantification of microtubule poleward flux speeds in RPE-1 PAGFP-αtubulin cells treated with indicated siRNAs. N = 3, n = 28-47 cells, ****p < 0.0001, one-way Anova test. **(L)** Percentage of RPE-1 GFP-Centrin1/GFP-CENPA cells with transient lagging chromosomes after indicated siRNA treatments. Lagging chromosomes were quantified as shown in (E); N = 6-7; n = 54-73 cells; *siCTRL vs siCAMSAP1* p = 0.0062, *siCTRL vs siWDR62* p = 0.0025, *siWDR62 vs siCAMSAP1+siWDR62* p = 0.0015, in one-way Anova test. All scale bars = 5 µm.

### Co-depletion of CAMSAP1 in WDR62-depleted cells suppresses activation of the 53BP1 and p21 and restores normal cell proliferation

Quantitative immunofluorescence indicated that CAMSAP1 co-depletion also suppressed the appearance of 53BP1 foci and the accumulation of nuclear p21 in WDR62-depleted cells (Figure 4A-C). Moreover, CAMSAP1 co-depletion also restored normal cell proliferation in a WDR62 siRNA background, as quantified with the 5-day CFSE assay (Figure 4D). This supported our hypothesis that reduced poleward microtubule flux rates and transient lagging chromosomes activate 53BP1 and drive p21 expression to induce a cell proliferation delay.

**Figure 4:**
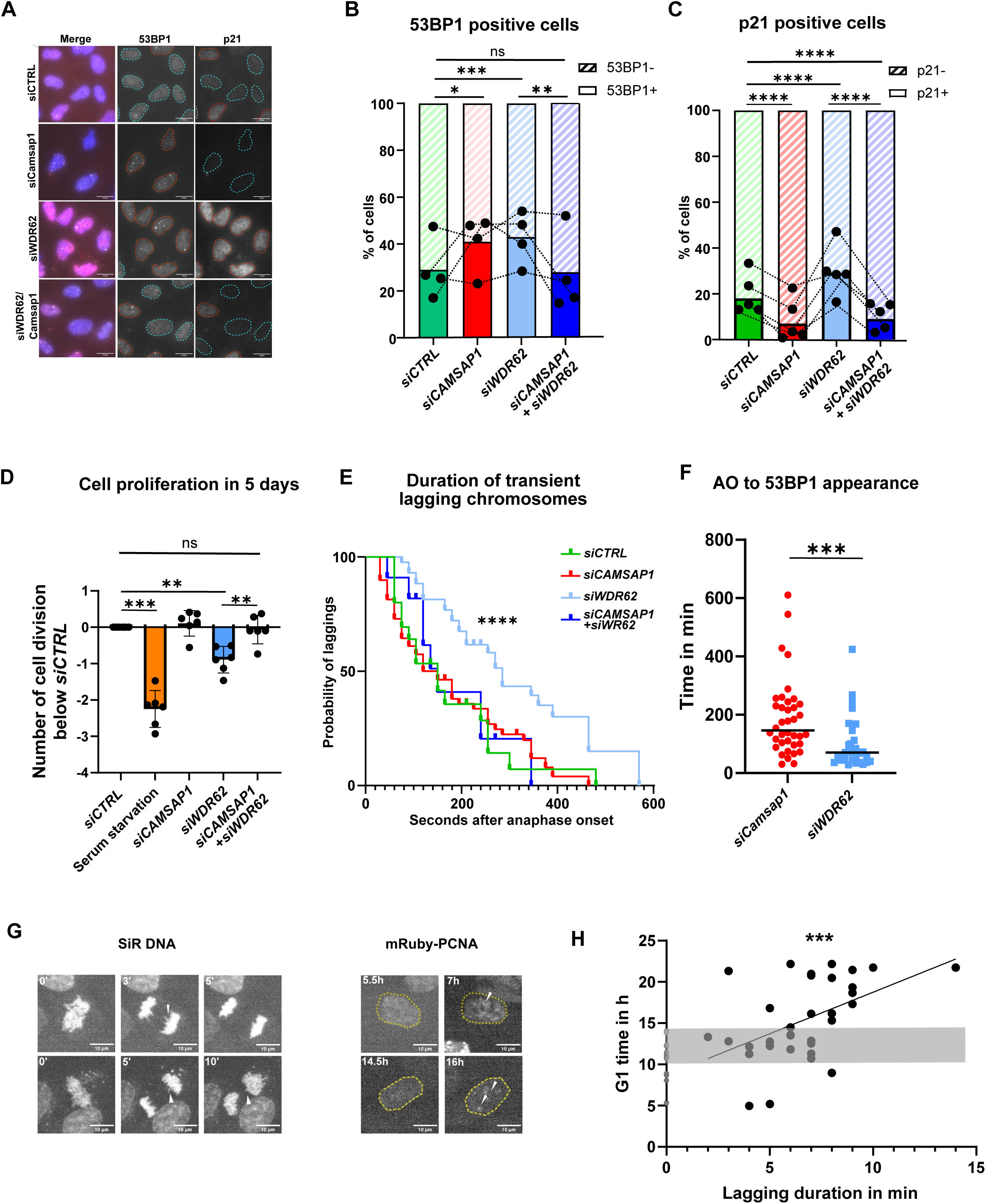
Camsap1 co-depletion rescues WDR62 RNAi phenotype. (**A**) Immunofluorescence images of interphase RPE1 cells treated with indicated treatments, stained with DAPI, 53BP1 and p21 antibodies. Blue dotted circles represent 53BP1-negative nuclei, red ones 53BP1-positive nuclei. Scale bars = 10 µm. (**B**) Quantification of 53BP1 positive cells after indicated siRNA treatments. Connected dots represent paired experiment. N = 4, n = 648 – 1440 cells; *siCTRL vs. siWDR62*, p = 0.0006; *siCTRL vs. siCAMSAP1*, p = 0.0381; *siWDR62 vs. siCAMSAP1 + siWDR62,* p = 0.0018, Fisher’s exact test with Bonferroni correction. (**C**) Quantification of p21 positive cells in indicated siRNA treatments. Connected dots represent paired experiment. Blue dotted circles represent p21-negative nuclei, red one p21-positive nuclei. N = 5, n = 1020 – 1330 cells; ****p < 0.0001; Fisher’s exact test with Bonferroni correction. (**D**) Quantification of cell divisions versus control depletion in CFSE assay in indicated treatments, N = 6, n = 69504-72579 cells; *siCtrl vs* serum starvation p = 0.0006, *siCTRL vs. siWDR62,* p = 0.0097: *siWDR62 vs. siCAMSAP1 + WDR62*, p = 0.0022, one-way Anova test; error bars represent SEM. (**E**) Quantification of persistence of transient lagging chromosomes in live RPE1 GFP-CenpA/GFP-Centrin1 cells after indicated treatments, N = 4, n = 43 - 59 cells; siCAMSAP1 vs siWDR62 ****p < 0.0001 in Mann-Whitney test. (**F**) Time of 53BP1 foci appearance in *siCAMSAP1* and *siWDR62*-treated RPE1 GRP-53BP1 live cells having experienced a lagging chromosome, N = 14, n =33-38 cells; ***p = 0.0001, Mann-Whitney test. (**G**) Live-cell imaging stills of WDR62-depleted mRuby-PCNA RPE1 cells stained with SiR-DNA; displayed is a cell with a rapidly resolved lagging chromosome (upper panels) or a slowly resolved lagging chromosome in anaphase (lower panel) with the corresponding panels a time point before and one after appearance of PCNA aggregates, indicative of G1/S transition (**H**) Scatter-plot of time of G1 duration vs persistence of transient lagging chromosomes, n= 49 cells; r = 0.6020, ***p = 0.0006, Spearman correlation test.

Nevertheless, we also noted that cells depleted of CAMSAP1 alone, neither accumulated p21 (Figure 4D) nor experienced a cell proliferation defect (Figure 4A), despite transient lagging chromosomes (Figure 3F) and an increased proportion of cells with 53BP1-foci (Figure 4B and C). How was this possible? Re-examining both live-cell GFP-CENP-A and GFP-53BP1 movies revealed key differences: (1) while transient lagging chromosomes in WDR62-depleted GFP-CENP-A cells persisted in anaphase for a median time of 3.5 mins, the corresponding median time in control-, CAMSAP1-, or CAMSAP1/WRD62-depleted cells was only 2 mins (p < 0.0001 in Mann-Whitney test); (2) 53BP1 foci appeared on average 1 hour after anaphase in WDR62-depleted cells, but only after 2 hours in CAMSAP1-depleted cells (p = 0.0001 in Mann Whitney test, Figure 4E and F). This suggested that the the probability to activate 53BP1 and delay cell proliferation might correlate with the persistence of transient lagging chromosomes. This would also be consistent with the fact that unperturbed cells often experience transient lagging chromosomes for brief periods without impairing cell proliferation. To test this idea, we recorded dual-colour live-cell SiR-DNA and mRuby-PCNA movies (a marker for the G1/S transition) at high temporal resolution and plotted at the single cell level the persistence of transient lagging chromosomes versus G1 duration (Figure 4G and H; note that in SiR-DNA-labelled cells transient lagging chromosomes are visible for longer times than in GFP-CENP-A cells). We found that daughter cells having experienced a transient lagging chromosome in anaphase had, as predicted, longer G1 (Fig. 4H; p = 0.0396 in Mann-Whitney test), and that their G1 duration correlated with the persistence of the transient lagging chromosomes in anaphase (r = 0.602, p = 0.0006 in Spearman correlation test). We conclude that poleward microtubule flux defects lead to transient lagging chromosomes that, if not rapidly resolved, will delay cell proliferation.

### Patronin co-depletion restores brain size and cognitive function in WDR62-depleted fly larvae

Given that primary microcephaly is caused by an exhaustion of neuronal progenitor cells, and that an extension of the cell cycle timing in murine progenitors causes a premature cell differentiation (*41*), we hypothesized that the proliferation impairment we observed, could explain the origin of microcephaly in organisms lacking WDR62. To test this hypothesis, we turned to *Drosophila melanogaster* larvae, where *WDR62* mutations cause a cell cycle delay and a brain size reduction (*42*). We used the UAS/Gal4 system (*43*) to express UAS-*WDR62* and UAS-*Patronin* RNAi constructs (*WDR62* RNAi and *Patronin* RNAi hereafter, Supplementary Figure 3B) to knock-down their respective mRNA specifically in the brain (*44*, *45*). We first tested whether Patronin co-depletion counteracts the effects of WDR62 RNAi on brain lobe size. To compare brain size of larvae of similar age, we staged egg-laying and stained the resulting third instar larvae with the DNA-marker DAPI and the neuroblast-specific polarity marker Miranda ((*46*); Figure 5A and supplementary Figure 3C). Consistent with previous studies, brain size (p < 0.0001) and neuroblast numbers in the central brain (p = 0.0041 in one-way Anova) was decreased by one quarter in *WDR62* RNAi compared to control conditions (Figure 5A-C). *PATRONIN* RNAi had no effect on its own but led to larger brain lobes and restored neuroblast numbers in a *WDR62* RNAi background (Figure 5A-C), indicating a rescue of the microcephaly phenotype. Miranda staining also allowed us to probe the polarity of the neuroblasts and spindle orientation vs the polarity axis ((*46*); supplementary Figure 3C and D). We observed the typical Miranda crescents in all conditions and found no change in spindle orientation (Figure 5D and supplementary Figure 3D), indicating that the brain size phenotypes occurred independently of cell polarity or spindle orientation.

**Figure 5:**
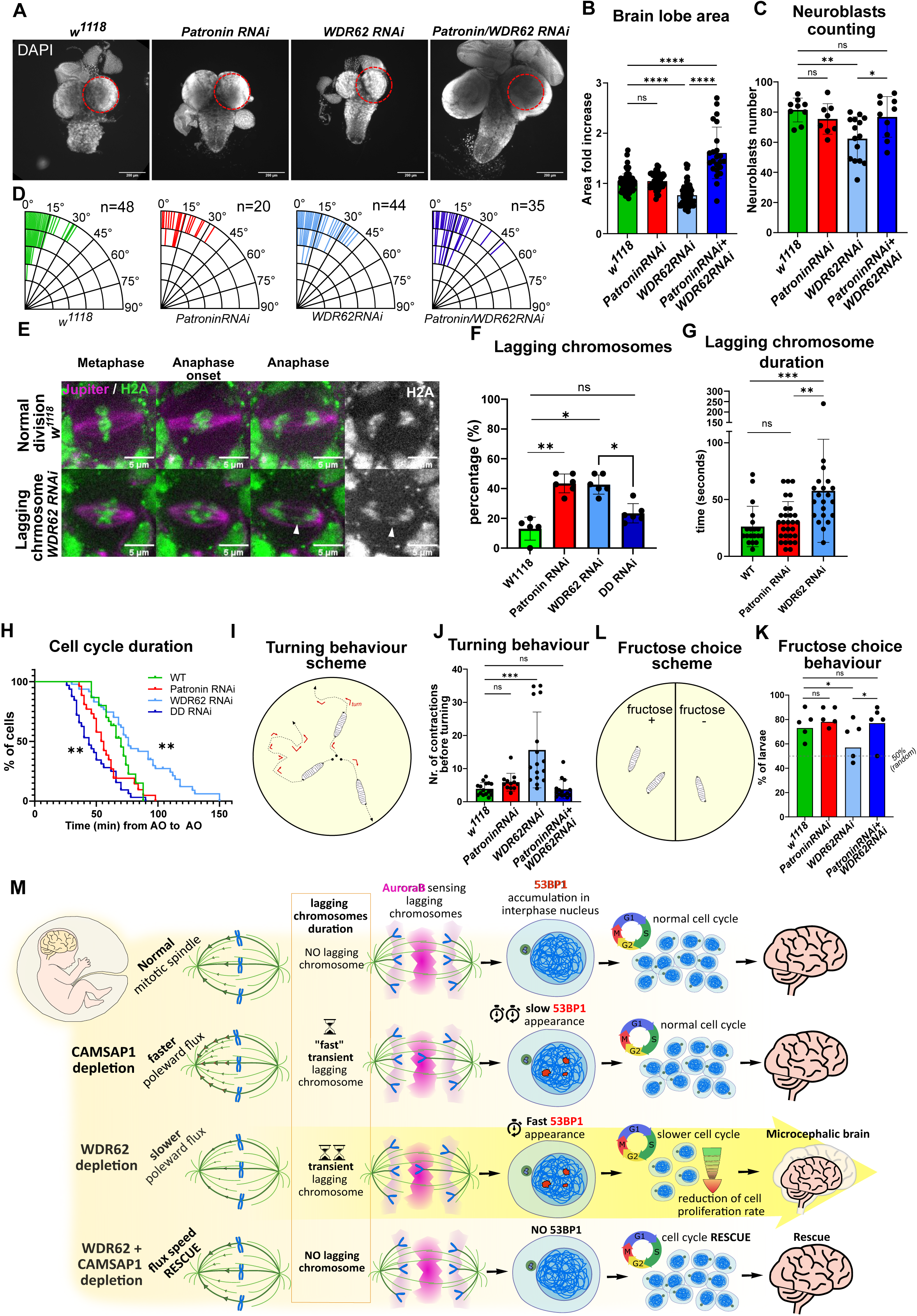
Co-depletion of Patronin rescues small brain phenotype in WDR62 RNAi background. (**A**) Brains extracted from *D. melanogaster* larvae of w^1118^, *PatroninRNAi*, *WDR62RNAi*, and *PatroninRNAi + WDR62RNAi*, stained with DAPI. The dotted red circle represents the area of the right lobe from w^1118^ brain superposed on *PatroninRNAi*, *WDR62RNAi*, *PatroninRNAi+ WDR62RNAi* brains. (**B**) Brain lobes area quantification of *D. melanogaster* larvae brains extracted from w^1118^ (n= 73 lobes), *PatroninRNAi* (n= 56), *WDR62RNAi* (n= 76), *PatroninRNAi+ WDR6RNAi* (n= 23) larvae., w^1118^ vs *WDR62RNAi* p < 0.0001, w^1118^ vs *PatroninRNAi + WDR62RNAi* p < 0.0001, *WDR62RNAi* vs *PatroninRNAi + WDR62RNAi* p < 0.0001, one-way Anova test. (**C**) Quantification of neuroblast stained with Miranda antibodies in brains extracted from w^1118^ (n= 9 brains), *PatroninRNAi* (n= 8), *WDR62RNAi* (n= 15), *PatroninRNAi+ WDR6RNAi* (n= 10) larvae, w^1118^ vs *WDR62RNAi* p = 0.0041, *WDR62RNAi* vs *PatroninRNAi + WDR62RNAi* p = 0.0324, one-way Anova test. (**D**) Quantification of the spindle orientation angles (as determined in Fig.5E) in neuroblasts from *D. melanogaster* larvae brain lobes from w^1118^ (n= 48 cells), *PatroninRNAi* (n= 20), *WDR62RNAi* (n= 44), *PatroninRNAi+ WDR6RNAi* (n= 35) larvae, all non-significant, Kruskal-Wallis test. (**E**) Live cell imaging of WT (upper panel) or *WDR62* RNAi neuroblasts in live larval brains in the Jupiter-Tomato/GFP-H2A strain. White arrow indicates lagging chromosome (**F**) Quantification of lagging chromosomes in indicated genetic background, n = 34-79 cells, W118 vs *PATRONIN RNAi* p < 0.0001, W118 vs *WDR62 RNAi* p < 0.0001, *WDR62 RNAi* vs *PATRONIN + WDR62 RNAi* p = 0.0004 in one-way Anova test (**G**) Quantification of the duration of lagging chromosomes in W118 (n = 19 cells), *PATRONIN RNAi (n = 29 cells)* and *WDR62 RNAi (n = 21 cells)* neuroblasts, W118 vs *WDR62 RNAi* p = 0.0004, *PATRONIN vs WDR62 RNAI* p = 0.0043, Kruskal-Wallis test (**H**) Quantification of cell cell cycle times in neuroblasts with indicated genetic background, n = 44 – 76 cells, W118 vs *WDR62 RNAi* p = 0.0096, W118 vs *PATRONIN+WDR62 RNAi* p = 0.0015, Log-rank Mantel-Cox test with Bonferroni correction (**I**) Scheme of larvae turning behaviour assay: the number of larva contraction before each change in direction was determined (see Supplementary Fig3E for examples of tracking). (**J**) Quantification of the contractions before change of direction in w^1118^ (n=16 larvae), *PatroninRNAi* (n= 12), *WDR62RNAi* (n= 16), *Patronin RNAi+ WDR6RNAi* (n= 15) larvae. w^1118^ vs *PatroninRNAi*, w^1118^ vs *WDR62 RNAi* p = 0.0001, *WDR62 RNAi* vs *PatroninRNAi + WDR62RNAi* p < 0.0001, Kruskal-Wallis test. (**K**) Scheme of larvae fructose choice assay: the proportion of larvae on the half containing fructose was counted after 5 minutes of free movements (See Supplementary Fig5D for examples of tracking). (**J**) Quantification of larvae number on the Fructose-containing half after 5 minutes of movements of w^1118^ (n=68 larvae), *PatroninRNAi* (n=58), *WDR62RNAi* (n=54), *PatroninRNAi+ WDR6RNAi* (n=53) larvae. w^1118^ vs *PatroninRNAi*, w^1118^ vs *WDR62RNAi* P = 0.0319, *WDR62RNAi* vs *PatroninRNAi + WDR62RNAi* P = 0.0390, Fisher’s exact test. (**M**) Proposed model for the origin of primary microcephaly origin: We postulate that cells can can detect lagging chromosomes via the Aurora B activity gradient. If these lagging chromosomes are only briefly in contact with Aurora B, as is the case in many wild-type or more frequently in CAMSAP1-depleted cells (fast poleward microtubule flux speed), this may lead to a late activation of 53BP1, but not an activation of p21. If these lagging chromosomes remain longer, as is the case in WDR62-depleted cells (slow poleward microtubule flux speed) this will lead to a rapid 53BP1 activation and p21 induction resulting in a cell cycle delay which can exhaust the neuroprogenitor cell pool. Re-equilibrating poleward microtubule flux rates by co-depleting CAMSAP1 and WDR62 prevents transient lagging chromosomes and rescues the primary microcephaly phenotype.

To test whether *WDR62, PATRONIN* and *WDR62 + PATRONIN* RNAi also led to transient lagging chromosomes in anaphase, we applied these depletions in a strain expressing the microtubule-marker Jupiter-GFP and the chromosome marker His2Av-mRFP (*47*, *48*), extracted the live brain from the larvae, and monitored neuroblast divisions by live cell imaging (Figure 5E). As in human cells, both *PATRONIN* and *WDR62* RNAi led to transient lagging chromosomes in anaphase, while their co-depletion suppressed this phenotype (Figure 5E and F). Similar to human cells, lagging chromosomes persisted twice as long in WDR62-depleted neuroblasts than in PATRONIN-depleted or WT cells (Figure 5F and G; p = 0.0043, Kruskal-Wallis test). Moreover, as RPE1 cells, WDR62-depleted neuroblasts had a normal mitotic timing (Supplementary Figure 3E), but a longer cell cycle (Figure 5H). In contrast, *WDR62 + PATRONIN RNAi* led to a shorter cell cycle, which could partially explain the resulting larger brain size (Figure 5H). We conclude that the depletions of PATRONIN and WDR62 in in situ neuroblasts fully phenocopy the orthologous depletion in human RPE1 cells.

At the functional level, to test whether the reduction in brain size in WDR62-depleted cells impairs larval cognitive functions, we performed two basic behavioural assays. In the first assay we recorded larvae on agar plates as they searched for food (Figure 5G and supplementary figure 3C). Wild-type larvae optimize their search strategy by turning by 90° after 3-4 contractions (*49*). In contrast, *WDR62* RNAi-treated larvae turned much less frequently, leading them to unidirectional movements (11 median contraction per turn, p < 0.0001; Figure 5H and supplementary Figure 3C). This defect was rescued by *Patronin* co-depletion (3 median contractions per turn; Figure 5H and supplementary Figure 3C). Second, we plated larvae on agar plates containing in one half fructose and in the other half no fructose and recorded the position of the larvae after 5 minutes. As previously described, wild-type larvae selectively chose the fructose containing half-plate (76%; (*50*)); in contrast, WDR62 RNAi larvae showed a much weaker preference for fructose (57% p = 0.0319 in Fischer’s exact test; Figure 5I and J, supplementary Figure 3D). This cognitive defect was again rescued by *Patronin* co-depletion (77%, Figure 5J). We concluded that co-depletion of Patronin rescues the microcephaly phenotype induced by WDR62 depletion in terms of brain size, neuroblast depletion and basic cognitive functions.

## DISCUSSION

Loss of function of WDR62 or ASPM causes over 80% of the primary microcephaly cases (*11*). These cases, however, are not due to the postulated cellular origins for primary microcephaly: excessive mitotic duration, loss or gain of chromosomes, or spindle orientation defects (*6*, *8–10*, *51*). Instead, our *in vitro* and *in vivo* results indicate that a deregulation of microtubule minus-end dynamics, which leads to transient lagging chromosomes that elicit an Aurora-B dependent activation of 53PB1 and the p53-target p21, impairs cell cycle progression thus limiting the number of neuronal progenitor cells (see model Figure 5K). We postulate that this response is due to a surveillance mechanism that reacts to chromosomes that are not segregated in time with the rest of the chromosome mass in anaphase, complementing the mitotic surveillance mechanism that checks whether cells have remained for too long in mitosis (*26–29*, *52*). Given that loss of many cell division genes associated to primary microcephaly leads to transient lagging chromosomes, we speculate that this mechanism could be a frequent cause of this disease.

One major difficulty when evaluating the potential cellular causes of primary microcephaly, has been the lack of effective rescue experiments that could validate the hypotheses. Indeed, it is difficult to correct for aneuploidy, spindle orientation defects or excessive mitotic timing without introducing additional mitotic defects that may have confounding effects. Here however, by combining WDR62 and CAMSAP1 depletion and restoring normal minus-end microtubule dynamics, we could rescue all the downstream effects of WDR62 depletion at the cellular level, establishing a causal chain of events. The fact that this co-depletion suppressed lagging chromosomes and the cell cycle delay in neuroblasts, while restoring their numbers and basic cognitive function in WDR62-depleted fly larvae, backs the hypothesis that defects in microtubule minus-end dynamics cause primary microcephaly in WDR62-depleted flies. Our data also imply that the speed of poleward microtubule flux needs to be balanced, as both an increase and a decrease in speed will lead to asynchronous anaphases, consistent with a recent study (*53*). Nevertheless, the rescue after WDR62/CAMSAP1 co-depletion is not perfect: co-depletion of *WDR62* and *Patronin* results in larvae with larger brain size. One potential cause could be the shorter neuroblast cell cycle we observed in this genetic background, which even at constant neuroblast numbers will generate more descendants. The neuroblasts in those larvae, however, did not resemble tumour cells, as their cell polarity was preserved.

Work in the last 15 years have shown that cells not only possess different mitotic surveillance mechanisms: a spindle checkpoint for kinetochore-microtubule attachment in metaphase (*54*), a surveillance mechanisms for aneuploidy (*55*), but also a mitotic duration surveillance mechanism, which blocks the cell cycle when cells spent more than 90 minutes in mitosis, whether they experience chromosome segregation errors or not (*26–29*, *52*). Here, we propose the existence of another, complementary surveillance mechanism that monitors chromosome movements in anaphase and activates 53BP1 in the presence of transient lagging chromosomes. This mechanism is distinct from the mitotic timing and aneuploidy surveillance systems, as WDR62-depleted cells have a normal mitotic timing and no increased aneuploidy. It allows cells to react to erroneous kinetochore-microtubule attachments that cannot be detected by the spindle checkpoint and do not extend mitotic timing, such as chromosomes with merotelic kinetochore-microtubule attachments, which often transiently lag in anaphase (one kinetochore bound by microtubules from both spindle poles; (*56*)). Previous studies had shown that the presence of an Aurora B gradient at spindle mid-zone can promotes their rapid correction and/or delay nuclear envelope reformation to prevent micronuclei formation (*23*, *57*, *58*). Here we postulate that the prolonged presence of in the high Aurora B activity zone also induces a long-term response, delaying cell proliferation. This response is linked to the proximity of anaphase chromatin to high Aurora-B activity and not the lagging per se, as it can be activated in Npl4- or UBASH3B-depleted cells without lagging chromosomes. Whether this 53BP1 activation depends on the ability of Aurora-B to delay nuclear envelope reformation (*35*, *36*), or its ability to phosphorylate chromatin proteins on lagging chromosomes (*37*), remains to be seen. The net result is that this response prevents the proliferation of cells having experienced such intermediate lagging chromosomes, which can lead to chromosome compaction and nuclear architecture defects in the absence of aneuploidy and impair post-natal growth in mice (*59*). We also postulate that this surveillance mechanism relies on a gradual response: chromosomes that lag only 1-2 minute at the level of kinetochores (as seen in GFP-CENP-A cells, Fig. 4E), or 5-7 minutes at the level of chromosome arms (as seen with SiR-DNA, Fig. 4H), such as in CAMSAP-1 depleted cells, do not impair cell proliferation; in contrast transient lagging chromosomes that persist for 3-4 minutes at the level of kinetochore or more than 7 minutes at the level of chromosome arms, as frequently seen in WDR62 depleted cells (Fig. 4E and H), are associated with a rapid 53BP1 foci formation, p21 activation, and a strong G1 delay. Such a semi-permissive mechanism also fits with the observation that short-lived transient lagging chromosomes can be observed in 20-30% of wild-type RPE1 cells.

Finally, we speculate that primary microcephaly might be caused by the frequent incapacity to rapidly satisfy this surveillance mechanism, due to transient lagging chromosomes. This disease is caused by the loss of over 30 different genes, many of which are linked to cell division and chromosome segregation. Although it is likely that several cellular mechanisms can contribute to neuronal progenitor exhaustion, we note that transient lagging chromosome are likely to appear in most of those genetic backgrounds, such as defective centriole biogenesis, impaired centrosome function, or loss of microtubule associated proteins, microtubule motors or kinetochore proteins, and might thus represent a common cellular origin. These mild defects would be sufficient to affect the expansion of neuronal progenitor cells, which have very short cell cycle, without impairing the overall development of the other organs. The mildness of these defects may also explain why patients having lost WDR62 or APSM do not display an elevated risk for cancer development, unlike patients with a predisposition to aneuploidy (*60*).

## ACKNOWLEDGEMENTS

We thank A. Khodjakov (State University of New York), R. Medema (Netherland Cancer Institute), E.Nagoshi (University of Geneva), A. Barr (Imperial College London) and R. Giet (University of Rennes) for reagents, the HumanaFly facility and M. Savitskiy (University of Geneva) for drosophila support, the Bioimaging Facility and N. Liaudet (University of Geneva) for microscopy support; the Flow Cytometry Core Facility (University of Geneva) for FACS support; S. Barbieri (University of Geneva) for help with data analysis, H. Maiato (University of Porto), Iva Tolic (Ruđer Bošković Institute), M. Gotta and her laboratory (University of Geneva), and members of the Meraldi laboratory for critical discussions. This work in the Meraldi laboratory is supported by the Swiss National Science Foundation project grants (208052 & 215124) and the University of Geneva.

## Author contributions

The project was initiated by E.D., D.I. and P.M. and directed by P.M.; E.D. and D.I. performed all experiments; A.T. contributed to the drosophila experiments; E.D., D.I. and P.M. analyzed and interpreted all the results; E.D., D.I., A.T. and P. M. wrote the manuscript.

## SUPPLEMENTARY MATERIAL

### MATERIALS AND METHODS

#### Cell culture and drug treatments

hTert-RPE1 (ATCC: CRL-4000), hTert-RPE1-53BP1-GFP (kind gift of R. Medema, Princess Maxima Center, Utrecht (*61*)), hTert-RPE1 GFP-Centrin1/GFP-CENPA (kind gift from A. Khodjakov, State University of New York), hTert-RPE1 mRuby-PCNA p21-GFP and hTert-RPE1^p21-/-^ (kind gifts of A. Barr, Imperial College London (*62*)) and hTert-RPE1 PAGFP-α-tubulin cell lines (*63*) were cultured in Dulbecco’s Modified Eagle’s Medium (DMEM; Thermo Fisher Scientific: 61965-026), supplemented with 10% FCS (Labforce, S181T), 100 U/ml penicillin and 100 mg/ml streptomycin (Life, 15140122). All cells were cultured at 37°C with 5% CO2 in a humidified incubator. For live-cell imaging, cells were cultured in eight-, four-, or two-well Ibidi chambers (Vitaris) in Leibovitz’s L-15 medium (Thermo Fisher Scientific, 21083027) supplemented with 10% FCS before imaging for maximum 12 h in the absence of CO_2_. The following drugs were added to the culture or imaging medium: 10 µM MG132 (Sigma Aldrich, C2211-5MG) for 30 min, 100 nM Doxorubicin (Sigma, D1515) for 12 h, 50 nM Aurora B inhibitor Barasertib (AZD1152, S1147, Selleckchem), 50 nM SiR-Tubulin (Spirochrome, SC002) for 2 h, 100 nM SiR DNA (Spirochrome, SC007) for 2 h, 100 nM SpY DNA (Spirochrome, SC101) for 5 h.

#### siRNA and plasmid transfections

For protein depletions, cells were transfected for 48 h in case of single depletions and 72 h in case of the double depletions with 20 nM siRNAs using Opti-MEM (Invitrogen, 31985-047) and Lipofectamine RNAiMAX (Invitrogen, 13778075) according to the manufacturer’s instructions. For the double depletions, a double amount of Lipofectamine RNAiMAX was used. The medium was replaced 24 h after transfection. The following sense strands of validated siRNA duplexes were used (all Qiagen unless indicated): siASPM (*16*), 5′-GCU AUA UGU CAG CGU ACU ATT-3′; siCamsap1 (validated in Supplementary Figure 3) 5′-CAUCGAGAAGCUUAACGAATT-3′; siWDR62 (Dharmacon; (*14*)), 5′-AGA CAA AGG UGA CGA GCA C-3′; siNPL4 (*39*), 5′-CAGCCUCCUCCAACAAAUCdTdT-3’; siUBASH3B – 1 (*40*), 5′-CCGCTTAAGGATGCTAACATT-3’, siUBASH3B - 2 (*40*),), 5′-AAGAGAGTTGTTCTTAGGTTA-3’ mixed 1:1 and as a negative control siCTRL (AllStars Negative Control siRNA; Qiagen, 1027281, Proprietary).

#### Immunofluorescence

To stain for WDR62, 53BP1, p21, yH2AX, Aurora B, NPL4, UBASH3B hTert-RPE1 cells were fixed at -20°C for 7 min with ice-cold methanol stored at -20°C. To stain for CAMSAP1, MCAK, hTert-RPE1 cells were fixed at room temperature for 15 minutes with a solution containing 4% Formaldehyde (Applichem, A3592), 2mM of PIPES (Applichem, A1079.0500), pH 6.8, 1mM of EGTA (Applichem, A0878.0100), and 0.2% Triton X-100 (Applichem, A4975.0100). To stain for α-tubulin cells were washed 30 seconds in a 37°C warm buffer containing 80mM M KOH-PIPES, 10mM MgCl2 (Applichem, 131396.1211), 5mM EGTA, 0.5% Triton-X and fixed at room temperature for 10 minutes in a 12.5% Glutaraldehyde (Sigma Aldrich, G5882), 80mM KOH-PIPES, 10mM MgCl2, 5mM EGTA and 0.5% Triton-X solution. Cells were washed for 7 minutes in a freshly prepared 0.1% NaBH4 (Sigma Aldrich, 16940-66-2) in PBS quenching solution. The samples were washed twice for 2 minutes in PBS. After all fixations cells were rinsed with PBS and blocked 1 hour in blocking buffer (PBS + 3% BSA (LabForce, S181T) and 1% N_3_Na (Applichem, A1430). After blocking, cells were washed thrice for 5 minutes with PBS and incubated 1 hour with the primary antibody diluted in blocking buffer. Next, cells were washed twice in PBS and incubated 1 hour with the secondary antibody diluted in blocking buffer. Coverslips were mounted on microscopes slides with DAPI Vectashield mounting medium (Vector Laboratories, H1200).

The following primary antibodies were used: recombinant human anti-α-tubulin (1:500; (*64*)), rabbit anti-CAMSAP1(1:1500; Novus Biologicals NBP1-26645), rabbit anti-WDR62 (1:1000; Bethyl A301-560A), rabbit anti-MCAK (1:1000; (*65*)), rabbit anti-53BP1 (1:1000; Cell Signalling Technology 4937), mouse anti-p21 (1:1000; Cell Signalling Technology 2947), mouse anti-yH2Ax (1:2000; EMD Millipore, 05-636), mouse anti-AuroraB (1:2000; BD Biosciences 611083), rabbit anti-NPL4(1:500; Novus Biologicals NBP1-82166), rabbit anti-UBASH3B (1:500; Proteintech 19563-1-AP). For secondary antibodies, Alexa Fluor–conjugated antibodies (1:400; Invitrogen) were used. The pictures used for the analysis were acquired using 60× and 100× (NA 1.4) oil objectives on Olympus DeltaVision wide-field microscope (GE Healthcare) equipped with a DAPI/FITC/TRITC/Cy5 filter set (Chroma Technology Corp.) and Coolsnap HQ2 CCD camera (Roper Scientific) running Softworx (GE Healthcare). Alternatively, they were also acquired using a HC PL APO CS2 63x/1.40 Oil objective on a Leica Stellaris 5 confocal microscope equipped with 405 nm, 488nm, 561nm and 638 nm lasers and two Hybrid S detectors (HyD S1 and HyD S2) running LAS X software (version: 4.5.0.25531).

#### Live-cell imaging

To monitor mitotic progression and chromosome segregation, hTert-RPE1 and hTert-RPE1-53BP1 GFP cells were seeded into Ibidi chambers (Vitaris) and treated with siRNAs as described above. Prior to imaging, cells were supplemented with L15 medium containing SiR DNA (Spirochrome #SC007) according to manufacturer protocole and recorded on a Nikon Eclipse Ti-E wide-field microscope (Nikon) equipped with a GFP/mCherry/Cy5 filter set (Chroma Technology Corp.), an Orca Flash 4.0 complementary metal-oxide-semiconductor camera (Hamamatsu), and an environmental chamber using NIS software (Nikon). For normal movies cells were imaged in the GFP and Cy5 channels for 12 h at 37°C, at 1-min 30 sec or 2 min intervals, in 17 steps of 1 µm Z-stacks, using a 60× (NA 1.51) oil objective and 2 × 2 binning. To correlate the persistence of lagging chromosomes (SiR-DNA) and the timing of appearance of GFP-53BP1 foci, a first movie was recorded in the Cy5 channel for 1 h at 37°C at 30 sec intervals, in 17 steps of 1 µm Z-stacks, using a 60× (NA 1.51) oil objective and 2 × 2 binning. The movie was stopped and restarted at the same positions using both the GFP and Cy5 channels for 12 h at 37°C at 3 min intervals, in 17 steps of 1 µm Z-stacks, using a 60× (NA 1.51) oil objective and 2 × 2 binning. For cell cycle duration measurement, hTert-RPE1 cells were seeded into Ibidi chambers and treated with siRNAs as described above. Prior to imaging, cells were supplemented with Leibovitz’s L-15 Medium and recorded on the same Nikon Eclipse Ti-E wide-field microscope using brightfield illumination for 30h at 37°C, at 3 min interval, in single plane, using a 10× objective and 2 × 2 binning. To correlate the persistence of lagging chromosomes (SiR-DNA) and the timing of G1 phase, hTert-RPE1 mRuby-PCNA p21-GFP were seeded into Ibidi chambers and treated with siWDR62 as indicated. Prior to imaging, cells were supplemented with Leibovitz’s L-15 Medium and SiR DNA. A first movie was recorded at 37°C using a ZEISS Lattice Lightsheet 7 microscope equipped with three laser lines (488 nm, 561 nm, and 640 nm), two Hamamatsu Fusion sCMOS cameras, and a water immersion 48x/1.0 NA detection objective. SiR DNA was recorded for 2 h at 1 min intervals, in 114 0.4 µm Z-stacks in the 640nm channel. The movie was stopped and restarted at the same positions using both the 561 (mRuby-PCNA) and the 640nm (SiR-DNA) for 20 h with 114 0.4 µm Z-stacks.

#### Anaphase kinetochore tracking assay

hTert-RPE1 GFP-centrin1/GFP-CENPA were treated with the reported siRNAs, and metaphase cells imaged for 15 min every 15 s using a 100× (NA 1.4) oil objective on an Olympus DeltaVision wide-field microscope (GE Healthcare) equipped with an environmental chamber maintained at 37°C and a GFP/RFP filter set (Chroma Technology Corp.). Z-stacks of 15-µm thickness with z-slices separated by 0.5 µm were imaged in the GFP channel with 2 × 2 binning to track kinetochores from metaphase until late anaphase. 4D images (XYZT) obtained were deconvolved in conservative mode and cropped using Softworx (GE Healthcare). The kinetochores and the poles were segmented using Imaris (Bitplane) during metaphase and anaphase. The kinetochores positions over time were analysed using a Matlab code ((*14*), https://github.com/AmandaGuerreiro/WDR62_2020).

#### Poleward microtubule flux measurement

hTert-RPE1 PA-GFP-α-tubulin cells were incubated with 10 µM MG132 for 30 minutes and imaged for 1 hour. Single focal planes of 150-nm pixel size were acquired using a 60× (NA 1.4) CFI Plan Apochromat oil objective on a Nikon A1r point scanning confocal microscope equipped with a 37°C heating chamber and running NIS elements software. A 3-pixel-thick and 100-pixel-long ROI parallel to the DNA and close to spindle centre was photoactivated with a 500-ms, 405-nm laser pulse at 50–100% intensity depending on the PA-GFP-α-tubulin expression levels and imaged every 20 seconds for 4 min in a single focal plane. The photoactivated mark was tracked manually for 120 s with Fiji (ImageJ). Distance between the photoactivated mark and the corresponding spindle pole was tracked in time and the displacement of the photoactivated mark over time was calculated.

#### Single cell sequencing

Cells were arrested in G1 phase using the CDK4/6 inhibitor Palbociclib (Cayman Chemical Company: 16273) at 100 nM for 12 hours. Cells were next sorted as single cells into a 384-well plate by FACS using the MoFlo Astrios cell sorter at the FACS facility of the Medical Faculty of the University of Geneva. At the single cell core facility of the ONCODE institute facility, nuclei in each well were digested with NlaIII, after which the genomic fragments (following end processing) were ligated to barcoded adapters containing a unique molecular identifier (UMI), cell-specific barcode, and T7 promoter allowing linear amplification by in vitro transcription (IVT). Libraries were sequenced on an Illumina Nextseq 500 at 16M reads. At the University of Geneva Bioinformatics facility, the raw fastq files were mapped to GRCH38 using the Burrows–Wheeler aligner. The mapped data were analyzed using custom scripts in R (available at GitHub: https://github.com/DIvanova-EDoria/master.git), which parsed for library barcodes, removed reads without a NlaIII sequence and removed PCR-duplicated reads. To perform quality control of the sequences, the following gates were set: minimum count: 10000 reads, maximum count: 75000 reads. The data which did not fail the QC were normalized to simulate the same amount of reads in every cell as following: 10000 * value / sum(value).

#### Metaphase spreads

Cells were incubated with 1mM Nocodazole (Sigma Aldrich, M1404-2MG) for 1h at 37C. Mitotic cells were shaken off, centrifuged and resuspended in 0.56% KCl (Applichem, A3582.0500) at room temperature for 6 min, before resuspending them dropwise in a 5 ml 3:1 MeOH/ CH3COOH (AppliChem, A2369) solution. The excess of supernatant was removed and 20ul of cells suspension was added dropwise on cooled microscopy slides (Epredia, 1.0mm) and left to dry in humid air chamber for 1h. The metaphase spreads were mounted using DAPI Vectashield and Epredia cover slips 22×22 mm.

#### CFSE labelling for cell proliferation assay

To measure cell proliferation over 5 days, hTert-RPE1, hTert-RPE1-PCNA-mRuby and hTert-RPE1-PCNA-mRuby p21 KO 2a (both kind gifts of Prof A. Barr, Imperial College London) cells were trypsinized, centrifuged, washed once with PBS, and resuspended in PBS at a concentration of 1 × 10^6^ cells/ml in the presence of 1 μM CFSE (BioLegend, 423801) for 20 min at 37 °C. The CFSE was quenched by adding 5 times the original staining volume of cell culture medium for 10 min. After a final washing step, a fraction of the cells was resuspended in PBS with 20mM Tris (Applichem, A1379.5000) and 0.1% FCS and quantified by flow cytometry for reference (see below). The remaining cells was seeded and treated with siRNAs for 5 days. After 5 days of incubation, CFSE signal intensity was measured using 488 nm laser on Beckman Coulter Cytoflex.

#### Image processing and 53BP1, p21 and **γ**H2AX quantifications

Immunofluorescence images were first converted to z-stack (sum slices) using ImageJ/Fiji and opened in QuPath 0.4.3 (*66*). To quantify 53BP1, p21 or γH2AX, nuclei were segmented using the DAPI channel. To detect 53BP1 foci, a code was used to determine foci with a minimal area of 0.2 μm^2^ and an intensity higher than the mean nuclear intensity plus 2 standard deviations. For p21 or γH2AX intensities, an automatized QuPath analysis was used to analyse p21 or γH2AX levels in segmented nuclei. The background was subtracted, and cells were counted as positive if their intensity was higher than the mean plus one standard deviation of the intensity in *siCTRL*-treated cells. Time-lapse movies were analysed manually with NIS Elements software to quantify mitotic timing and presence of 53BP1 foci, taking into account cells 4h after mitosis. Time-lapse movies were also analysed manually with NIS Elements software to quantify the presence or absence of transient lagging chromosomes. These were quantified as present when DNA clouds in anaphase were > 6um apart and the lagging was > 2um long. For siRNA immunofluorescence quantification, 3D images were analysed manually with ImageJ/Fiji. Aurora B intensity in anaphase was quantified using surface tool on DNA signal Imaris (Bitplane) 10.

All the data were plotted using GraphPad Prism 8.2.1.

### D. melanogaster strains

The following sources of different D. melanogaster lines were used in this study:

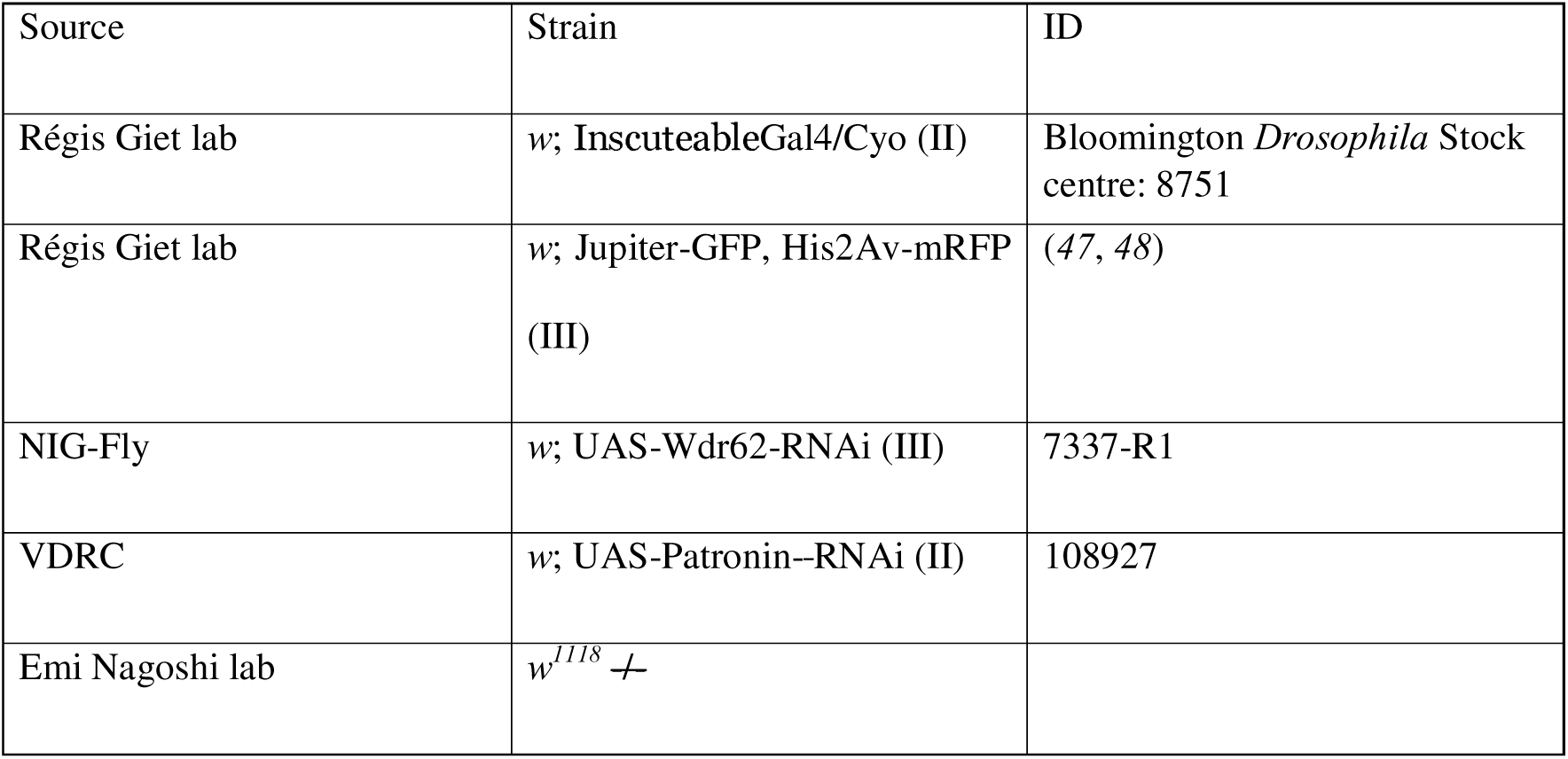

| Source | Strain | ID |
| --- | --- | --- |
| Régis Giet lab | *w*; InscuteableGal4/Cyo (II) | Bloomington *Drosophila* Stock centre: 8751 |
| Régis Giet lab | *w*; Jupiter-GFP, His2Av-mRFP (III) | (47, 48) |
| NIG-Fly | *w*; UAS-Wdr62-RNAi (III) | 7337-R1 |
| VDRC | *w*; UAS-Patronin-RNAi (II) | 108927 |
| Emi Nagoshi lab | *w*<sup>1118</sup> |  |

*D. melanogaster* fruit flies were raised at 25 °C in standard medium.

### Immunofluorescence of *D. melanogaster* brain larvae

3rd instar larvae were collected 120h after an egg laying of 5h. Larval brains were dissected in testis buffer (TB, 183 mM KCl, 47 mM NaCl, 10 mM Tris, and 1 mM EDTA, pH 6.8 MT) and brains were fixed at room temperature for 20 min in TB + 10% FA and 0.01% Triton-X. Brains were washed 2 times x 15 minutes in PBS and then 2 times x 15 minutes in PBS + 0.1% Triton-X. Brains were blocked for 1 hour in PBSTB (1% BSA, 0.1% Triton-X in PBS). Brains were incubated with the primary antibodies in PBSTB for 1 hour at room temperature. After 1 wash with PBS, brains were incubated with secondary antibodies in PBSTB for 1 hour at room temperature. After washes, samples were mounted on microscope slides with DAPI Vectashield mounting medium (to avoid brain deformation, for brain size measurements, spacer were used while mounting microscope slides).

The following primary antibody was used: rat anti-Miranda (1:500, Abcam, ab197788). For secondary antibodies, Alexa Fluor–conjugated antibodies (1:400; Invitrogen) were used. For neuroblast counting and spindle orientation experiments, images were acquired using a HC PL APO CS2 40x/1.40 air objective on a Leica stellaris 5 confocal microscope equipped with 405 nm, 488nm, 561nm and 638 nm lasers and two Hybrid S detectors (HyD S1 and HyD S2) running LAS X software (version: 4.5.0.25531). For brain size image acquisitions, a Nikon Eclipse Ti-E wide-field microscope (Nikon) equipped with a GFP/mCherry filter set (Chroma Technology Corp.), an Orca Flash 4.0 complementary metal-oxide-semiconductor camera (Hamamatsu), and an environmental chamber using NIS software was used.

To measure brain lobe sizes, the average diameter of one brain lobe was calculated as the mean of two measurements in orthogonal orientation (length + width / 2) using Fiji. The brain lobe shape was approximated to a circle shape and the area was calculated with the formula: Area=(π x diameter^2^)/4

The number of neuroblasts in the central brain was calculated as the number of Miranda positive cells, manually using FIJI, and the average of the two lobes was reported. For spindle orientation, Fiji from ImageJ software was used to calculate the angle between the spindle poles axis and a line perpendicular to Miranda signal.

### Live cell imaging of *D. melanogaster* brain larvae

3rd instar larvae were collected 120h after an egg laying of 5h. Larval brains were dissected in Schneider’s Drosophila Medium (Gibco, 21720024) and transferred in testis buffer into Ibidi chambers (Vitaris). Brains were then imaged on Nipkow spinning disk microscope equiped with EC Plan-Neofluar 10x / 0.3 Ph1 M27 WD=5.2mm and LCI Plan-Neoflur 63x / 1.3 Imm Korr DIC for Water Silicone Glycerole immersion WD=0.15-0.17 objectives and 405 nm, 488nm, 561nm and 638 nm lasers. For transient lagging chromosome and mitotic timing experiment, a 63x glycerol objective was used; metaphase cells were selected and recorded every 6 sec using 488nm and 561nm lasers until they were in late anaphase. QC gate of 1µm was used to differentiate a transient lagging chromosome. For cell cycle duration, a 10x objective was used and brains were acquired using 488nm and 561nm lasers every 2 min with a z-step of 1µm for a total of 31 steps.

#### Cognitive tests

For the running assay, all 3^rd^ instar larvae were recorded directly after collection on a 0.8% agar 10 cm (500 ml water, 4 g agar (Roth, Karlsruhe)) surface. Prior to recording, larvae were briefly washed with PBS. Turning frequency is indicated as the number of peristaltic movements a larva needs to make before turning. Movies of 5 min were made and analysed by hand selecting a 1 min interval. For fructose assay, all 3^rd^ instar larvae were recorded directly after collection on a 0.8% agar 10 cm (500 ml water, 4 g agar (Roth, Karlsruhe)) surface, containing in one half 0.2M Fructose (Sigma Aldrich, F0127, 3.6g in 100 ml 0.8% agar) and no fructose in the other half. Prior to recording, larvae were briefly washed with PBS. Fructose preference was evaluated by setting the larvae in the middle of the dish and allowing them to choose a side. The movements were recorded, and the final choice was analysed by the end of 5 min.

## SUPPLEMENTARY FIGURE LEGENDS

**Supplementary Figure 1:**
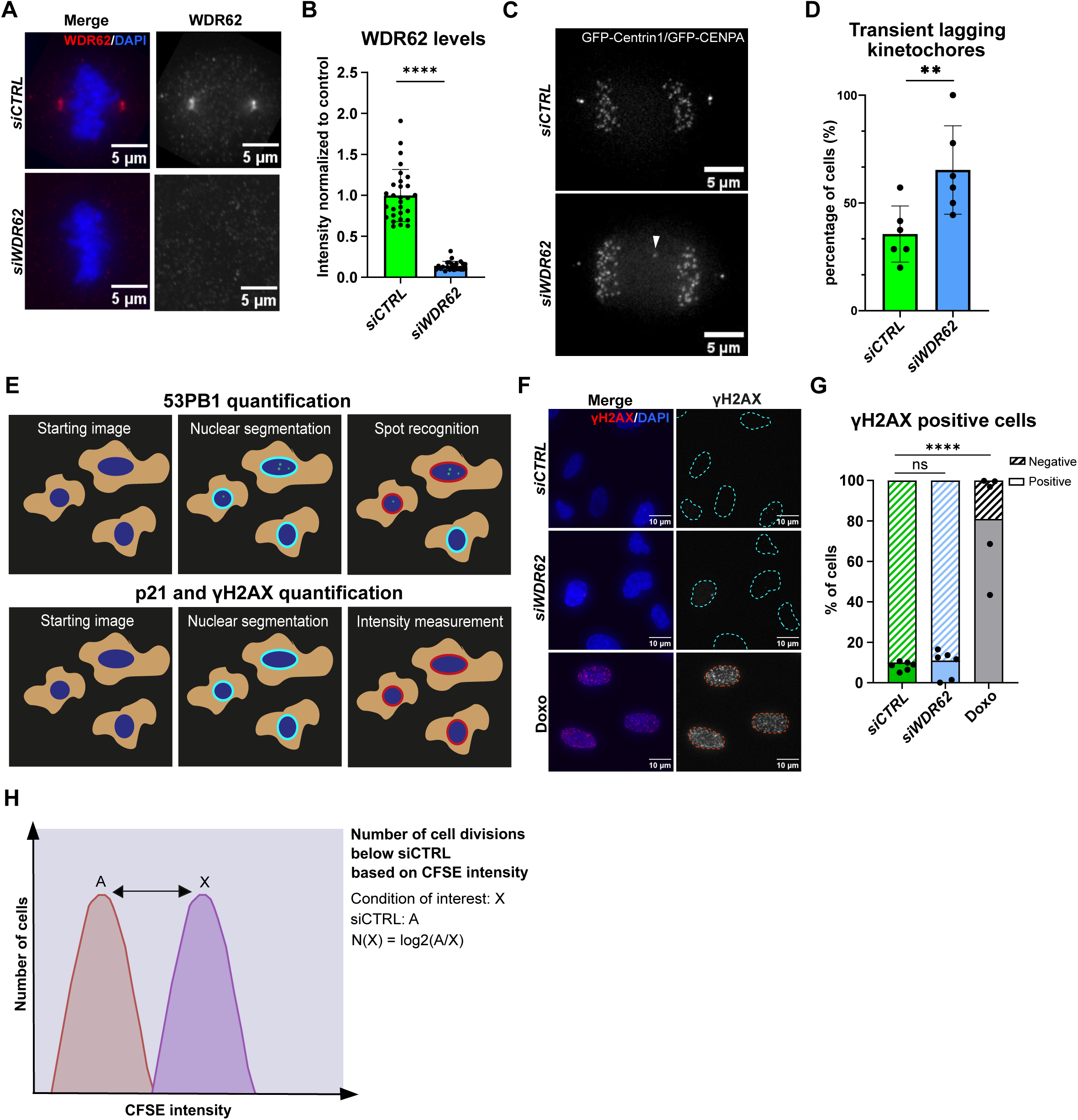
WDR62 depletion, and quantification of γΗ2AX, 53BP1 and p21. (**A**) Immunofluorescence images of metaphase RPE1 cells treated with *siCTRL* and *siWDR62* and stained with DAPI and WDR62 antibodies. Scale bars = 5 µm. (**B**) Quantification of WDR62 levels in RPE1 cells treated with *siCTRL* or *siWDR62* cells normalized to *siCTRL* values. N = 4 independent experiments, n = 49-81 cells, ****, p < 0.0001, unpaired t-test. Error bars represent SEM. (**C**) Live stills of GFP-centrin1/GFP-CENP-A RPE1 anaphase cells treated with *siCTRL* or *siWDR62*. White arrow indicates transient lagging chromosome in *siWDR62*-treated cell. (**D**) Quantification of GFP-Centrin1/GFP-CENP-A RPE1 cells with transient lagging chromosomes in anaphase after *siCTRL or siWDR62* treatment. N= 6; n = 54-55 cells; \**\** p = 0.0015 in paired t-test. Note that the data represented here are a duplicate of the quantification in Figure 3L. (**E**) Schematic representation of automated 53BP1 (upper panel) and p21/yH2AX analysis (lower panel). Nuclei were segmented using DAPI staining, and, for 53BP1, foci were counted as positive if the spots displayed an intensity than was higher than the nuclear mean plus 2 standard deviations; for p21/yH2AX the intensity of nuclei in corresponding channel were measured and counted as positive if their intensity was higher than the mean plus one standard deviation in *siCTRL*-treated cells. (**F**) Immunofluorescence images of interphase RPE1 cells, treated with *siCTRL*, *siWDR62* or 100 nM Doxorubicin, stained with DAPI and γH2AX antibody. Blue dotted circles represent γH2AX-negative nuclei, red ones γH2AX-positive nuclei. Scale bars = 10 µm. (**G**) Quantification of yH2AX positive cells after *siCTRL, siWDR62* or Doxorubicin treatment. N = 6, n = 834 – 2234 cells; *siCTRL* vs. Doxorubicin, ****, p < 0.0001, Fisher’s exact test with Bonferroni correction. (**H**) Schematic representation the Carboxyfluorescein succinimidyl ester (CFSE) assay: CFSE is a cell-permeable dye that covalently couples to intramolecular molecules via its succinimidyl group; since after a CFSE pulse its intensity decreases by half after each cell division, it can be used as a pulse-chase assay to quantify cell proliferation. In all our assays we compared the indicated siRNA-treatments to control depletion. To calculate the number of cell divisions above serum starvation we determined by FACS the peak intensity of a control-depleted cells (A) and divided it by the peak intensity of the condition of interest (X). The log2 of (A/X) indicates the number of divisions cells have undergone in comparison to control-depleted cells.

**Supplementary Figure 2:**
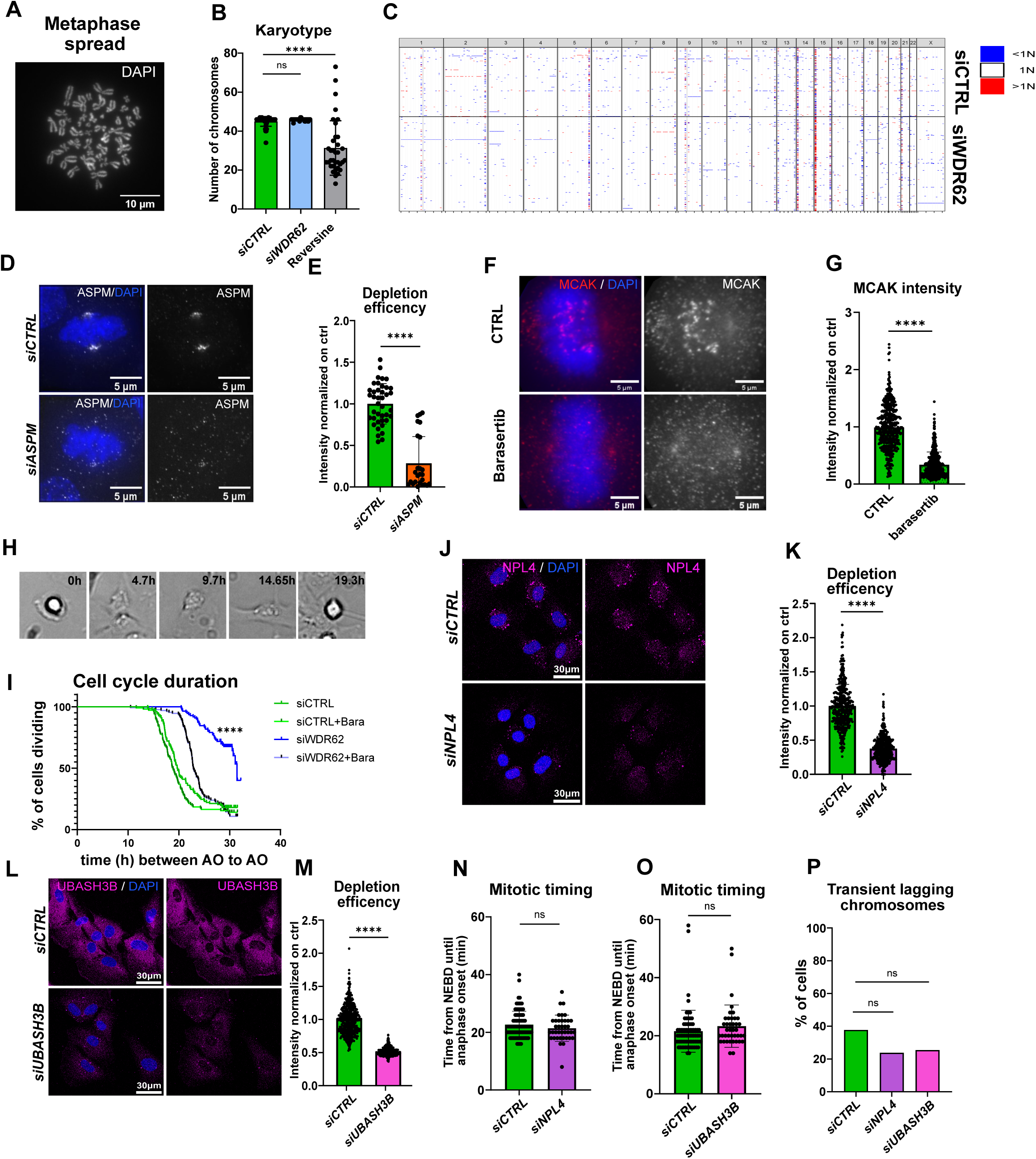
ASPM, NPL4, UBASH3B siRNA validation and characterization. (**A**) Chromosomes spread of RPE-1 cells stained with DAPI. Scale bar = 10 µm (**B**) Quantification of chromosomes number of RPE-1 cells treated with *siCTRL*, *siWDR62* or Reversine. N = 7; n = 35-42 cells; *siCTRL* vs Reversine ****p < 0.0001, Kolgomornov-Smirnov test with Bonferroni correction (**C**) Heat map of chromosome parts copy number variations across 384 control-depleted or 384 WDR62-depleted RPE1 cells (**D**) Immunofluorescence images of RPE-1 cells treated with *siCTRL* or *siASPM*, and stained for ASPM and DAPI. Scale bars = 5 µm (**E**) Quantification of ASPM levels in RPE-1 cells treated with *siCTRL* and *siASPM*. ASPM intensity was normalized to *siCTRL* in each replicate. N = 3, n = 26-40 cells; p= 0.0230, student t-test. (**F**) Immunofluorescence image of RPE-1 cells treated with DMSO (CTRL), Barasetrib and stained for MCAK and DAPI. Scale bars = 5 µm (**G**) Quantification of MCAK levels on kinetochores in RPE-1 cells treated with DMSO (CTRL) and Barasertib. MCAK intensity was measured on 10 kinetochores randomly selected per cell, MCAK values were normalized on the average intensity of the CTRL in each replicate. N = 3, n = 438-458 kinetochores, p = 0.0022, student t-test. (**H**) Phase-constrast live cell movies of RPE1 cells, illustrating how the cell cycle duration was determined using the time period between two anaphase onsets (**I**) Quantification of cell cycle time in the overall pool of cells subject to the indicated treatments. N = 3, n = 110-129 cells, ****p < 0.0001, Log-rank Mantel-Cox test with Bonferroni correction (**J**) Immunofluorescence images of RPE-1 cells treated with *siCTRL* or *siNPL4* stained for NPL4 and DAPI. Scale bars = 5 µm (**K**) Quantification of NPL4 levels in RPE-1 cells treated with *siCTRL* or *siNPL4*. NPL4 intensity was normalized to *siCTRL* in each replicate. N = 3, n = 419-430 cells; p = 0.0002, student t-test. (**L**) Immunofluorescence images of RPE-1 cells treated with *siCTRL* or *siUBASH3B* stained for UBASH3B and DAPI. Scale bars = 5 µm (**M**) Quantification of UBASH3B levels in RPE-1 cells treated with *siCTRL* or *siUBASH3B*. UBASH3B intensity was normalized to *siCTRL* in each replicate. N =3, n = 429-442 cells; ****p < 0.0001, student t-test. (**N** and **O**) Quantification of mitotic timing of RPE-1 cells treated with *siCTRL*, (N) *siNPL4* or (O) *siUBASH3B*, based on SiR-DNA live cell movies. Mitotic time is calculated from nuclear envelope break down till anaphase onset. N = 3, n = 39-90 cells. ns in student t-test (**P**) Quantification of transient lagging chromosomes in live PRE-1 cells stained with SiR-DNA and treated with *siCTRL*, *siNPL4* or *siUBASH3B*. ns in one-way ANOVA test.

**Supplementary Figure 3:**
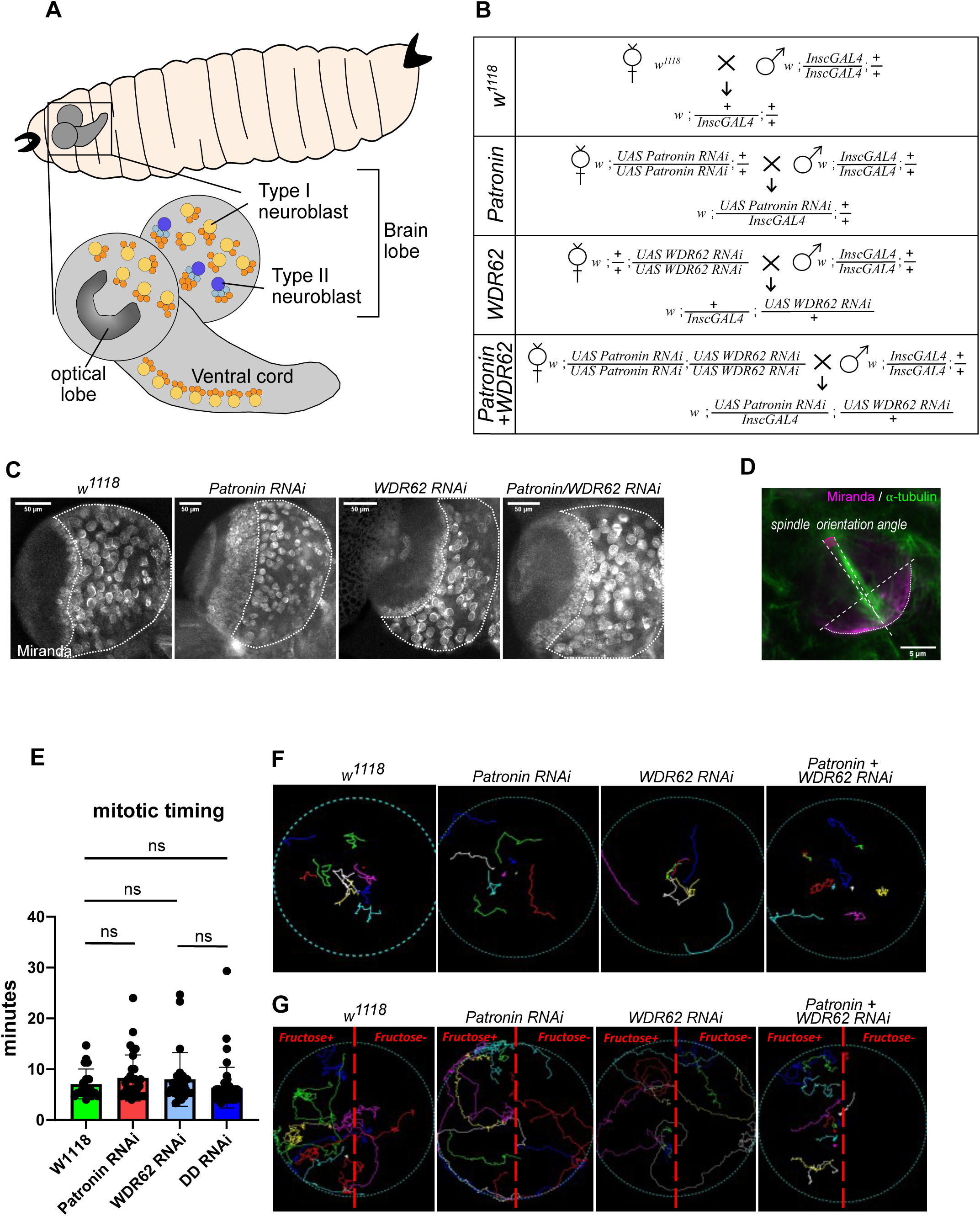
**(A)** *D. melanogaster* larva brain scheme, **(B)** *D. melanogaster* crosses used for the experiments reported in Figure 5. Wild-type (w^1118^), *PatroninRNAi*, *WDR62RNAi*, *PatroninRNAi+ WDR6RNAi* flies were crossed with *InscGAL4* flies in order to obtain larvae expressing the RNAi in type I and type II neuroblasts. (**C**) Extracted *D. melanogaster* larvae brains from w^1118^, *PatroninRNAi*, *WDR62RNAi*, *PatroninRNAi+ WDR6RNAi* and stained for Miranda and DAPI. The dotted white line highlights the shape of lobes. (**D**) Immunofluorescence from a larva brain stained for Miranda and α-tubulin. Dotted lines represent the pole-to-pole axe and the cellular axe perpendicular to the Miranda signal (spindle orientation angle). The magenta line represents the angle between the 2 axes. (**E**) Quantification of mitotic time (nuclear envelope breakdown till anaphase) in neuroblasts of indicated genetic background. Errors bars indicate SEM. (**F and G**) Examples of tracking of w^1118^, *PatroninRNAi*, *WDR62RNAi*, *PatroninRNAi+ WDR6RNAi* larvae movements on agar plates. Each colour is a single larva tracking. In (D) the agar plates are composed of a half agar Fructose+ and the other half agar Fructose-.

